# Glycan shield of the ebolavirus envelope glycoprotein GP

**DOI:** 10.1101/2022.02.07.479410

**Authors:** W. Peng, V. Rayaprolu, A.D. Parvate, M.F. Pronker, S. Hui, D. Parekh, K. Shaffer, X. Yu, E.O. Saphire, J. Snijder

## Abstract

The envelope glycoprotein GP of the ebolaviruses is essential for host cell attachment and entry. It is also the primary target of the protective and neutralizing antibody response in both natural infection and vaccination. GP is heavily glycosylated with up to 17 predicted N-linked sites, numerous O-linked glycans in its disordered mucin-like domain (MLD), and three predicted C-linked mannosylation sites. Glycosylation of GP is important for host cell attachment to cell-surface lectins, as well as GP stability and fusion activity. Moreover, it has been shown to shield GP from neutralizing activity of serum antibodies. Here, we use mass spectrometry-based glycoproteomics to profile the site-specific glycosylation patterns of ebolavirus GP. We detect up to 16 unique O-linked glycosylation sites in the mucin-like domain, as well as two O-linked sites in the head and glycan cap domains of the receptor-binding GP1 subunit. Multiple O-linked glycans are observed at the S/T residues of N-linked glycosylation sequons, suggesting possible crosstalk between the two types of modifications. We also confirmed the presence of C-mannosylation at W288 in the context of trimeric GP. We find heterogenous, complex N-linked glycosylation at the majority of predicted sites as expected. By contrast, the two conserved sites N257 and N563 are enriched in unprocessed high-mannose and hybrid glycans, suggesting a role in host-cell attachment via DC-SIGN/L-SIGN. We discuss our findings in the context of antibody recognition to show how glycans contribute to and restrict neutralization epitopes. This information on how N-, O-, and C-linked glycans together build the heterogeneous glycan shield of GP can guide future immunological studies and functional interpretation of ebolavirus GP-antibody interactions.

## Introduction

Ebola virus is a member of the *Filoviridae* family [1, 2]. Since its initial discovery in 1976, it has caused recurring outbreaks of disease in Central and West Africa upon spillover into the human population from an as-yet unidentified animal host reservoir, or recrudescence from convalescent humans [3, 4]. Detection of viral RNA and isolation of a new ebolavirus species (Bombali) from bats have pointed to these animals as a likely reservoir [3–5], similar to the related Marburg virus (MARV) for which the evidence is more established [6–8]. Outbreaks of ebolaviruses have typically been limited to the order of 10-1000 cases by contact tracing and isolation, but in 2013-2015 an outbreak with over 28000 confirmed cases and over 11000 deaths occurred in Sierra Leone, Liberia and Guinea [2, 9]. This outbreak accelerated the development of an effective vaccine and improved therapies against Ebola virus disease [10]. Still, clinical manifestation of Ebola virus infection has historically been associated with mortality rates ranging from 30% to 90% and even the most successful therapies to date provide only a modest improvement of mortality rates and don’t offer a cure for advanced disease [2, 10]. Six species of ebolavirus have currently been discovered, including Ebola (a.k.a. Zaire; EBOV), Sudan (SUDV), Bundibugyo (BDBV), Tai Forest (TAFV), Reston (RESTV) and Bombali (BOMV), of which all but the latter two are known to cause severe disease in humans [2].

The ebolaviruses are enveloped and contain an 18kb genome of non-segmented, negative sense, single-stranded RNA that encodes seven genes: *NP, VP35, VP40, GP, VP30, VP24* and *L* [2]. The *GP* gene encodes the full-length envelope glycoprotein (GP) as well as two truncated secreted versions (sGP and ssGP) by transcriptional editing [11–13]. The full-length envelope GP is a trimeric class I viral fusion protein and plays an important role in host cell attachment and entry [11]. Following virus internalization by macropinocytosis [14–17], GP binds the Niemann-Pick C1 (NPC1) receptor, triggering fusion of the viral envelope with the host membrane, thereby delivering the ribonucleoprotein complexes in the cytosol where replication will take place [18–20]. GP is also the primary target of antibodies produced upon natural infection or vaccination and of monoclonal antibodies developed as antiviral therapeutics [21]. Full-length GP is translated as a ~670 amino acid precursor and is cleaved by host furin into two disulfide-linked subunits: GP1 and GP2 [22, 23]. GP1 is responsible for receptor binding and consists of 4 domains: base, head, glycan cap and mucin-like domain (MLD) [11]. The GP2 subunit contains the fusion peptide and has the strongest conservation between different members of the filoviruses [24–26].

There are up to 17 N-linked glycosylation sites in ebolavirus GP, 15 of which are in the GP1 subunit, primarily in the glycan cap and MLD (see Figure 1A). The N-linked glycans mediate host-cell attachment through C-type lectins DC-SIGN/L-SIGN [27–29] and have been implicated in shielding GP from binding by neutralizing antibodies [28, 30–36]. Whereas the overall sequences and especially the N-linked glycosylation sites in the base, head and glycan cap are relatively well conserved among ebolavirus species, those in the MLD are highly variable. Further, the MLD is also modified with numerous O-linked glycans. The GP2 subunit contains two N-linked glycosylation sites that are conserved in all known mammalian filoviruses and play important roles in GP expression, stability and cell entry [36, 37]. Besides the numerous N- and O-linked glycans, there are also two predicted tryptophan C-mannosylation motifs in GP. These motifs consist of a WXXW sequence near the glycan cap, and a tandem WXXWXXW sequence in the membrane-proximal region of GP, where a mannose residue may be linked to the C2 atom of the first tryptophan’s indole group. The biological function of C-mannosylation is generally not well understood, but known to play a role in the folding, stability and trafficking of secreted glycoproteins, including components of the complement system and gel-forming mucins [38, 39]. The C-mannosylation motifs in GP are conserved in all ebolavirus species and the related Lloviu virus (LLOV), but not Marburg virus (MARV). So far, C-mannosylation in the glycan cap has been confirmed in the secreted version sGP [40], but its presence in full-length GP and role in the infection cycle remain unclear.

**Figure 1.**
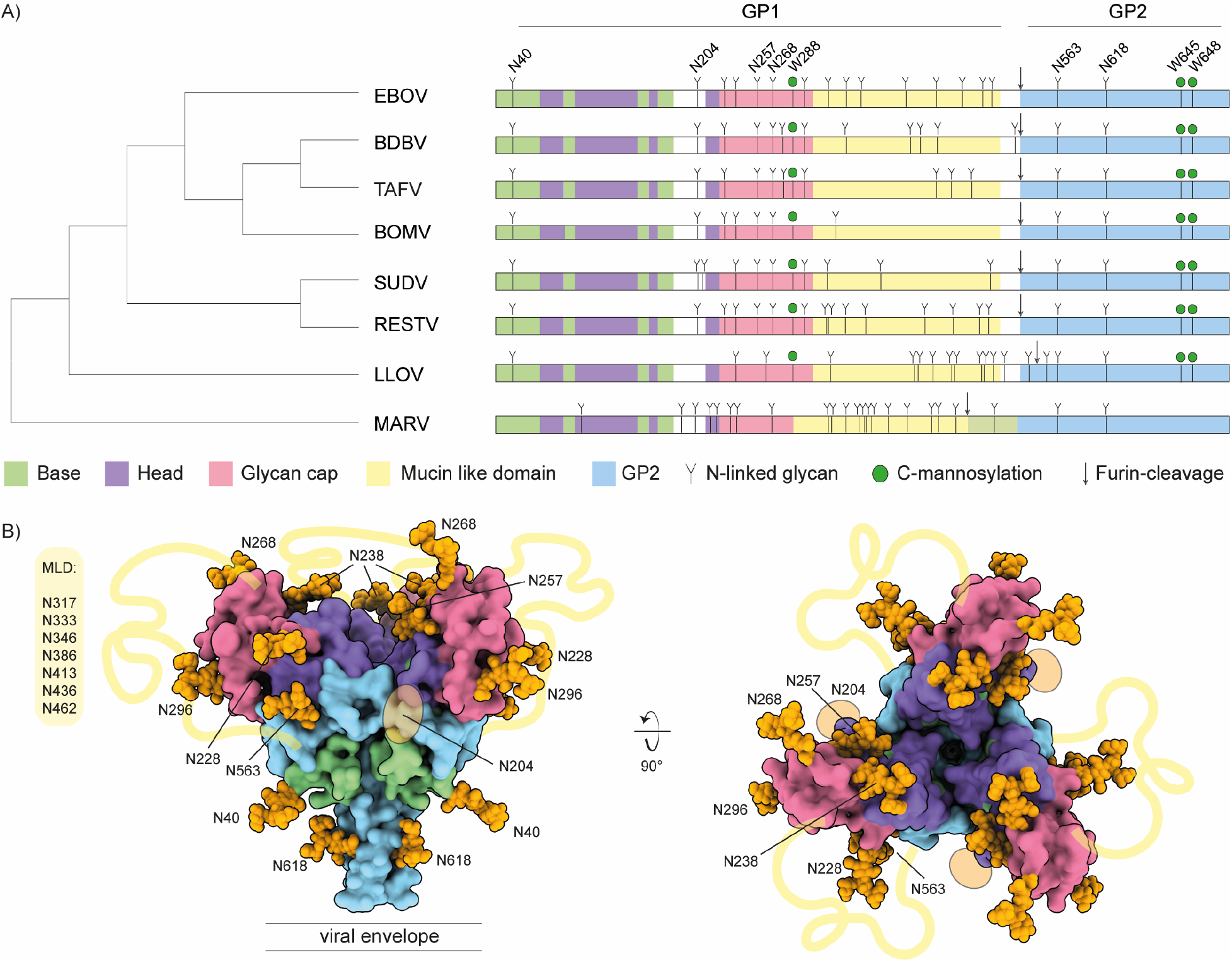
A) Schematic of filovirus GP domain structure with annotated N-linked glycosylation and C-mannosylation. B) Pseudomodel of EBOV GP with core pentasaccharide of N-linked glycans shown as orange spheres.

Glycomics studies have confirmed the presence of both complex N- and O-linked glycans in GP [30, 35], but little is known about the site-specific patterns of glycan processing. As glycans play a crucial role in host cell attachment and immune evasion, a better understanding of these patterns in the context of GP structure may help understand mechanisms of infectivity and epitope shielding. Here, we present an in-depth glycoproteomics study of the N-, O- and C-linked glycans of ebolavirus GP. We compare recombinant soluble GP ectodomains of EBOV and BDBV from both human HEK293 and insect S2 cells (material from S2 cells was included as it is often used as a diagnostic and research tool). We demonstrate that the conserved N-linked glycans at N257 and N563 are enriched in under-processed high-mannose and hybrid structures in both viral species and cellular expression platforms, suggesting a specific role in host cell attachment through binding of the cell surface lectins DC-SIGN/L-SIGN (which have a markedly higher affinity for high-mannose glycans). We observe that the MLD is modified by numerous O-glycans, comprising a mixture of truncated Tn-antigen and extended, sialylated core 1 and 2 structures, depending on the expression platform. Moreover, we find several O-linked glycosylation sites within the serine/threonine residues of N-linked glycosylation sequons, as well as evidence for O-linked glycans outside the MLD in both EBOV and BDBV GP. We also confirm C-mannosylation in the glycan cap of both ebolavirus species, which only occurs in the HEK293 expression platform. These key findings were confirmed in glycoproteomics experiments on virus-like particles formed by co-expression of full-length GP and VP40. We discuss the observed glycosylation profile in the context of known structures of GP in complex with neutralizing antibodies. Our findings provide a framework to understand the contributions and restrictions of GP glycosylation to the neutralization epitopes of antiviral antibodies.

## Results

We compared the pattern of predicted N-linked glycosylation sites (NXS/T) and C-mannosylation sites (WXXW) of all known ebolavirus species and the related filoviruses MARV and LLOV (see Figure 1A and the sequence alignment in Supplementary Figure S1). The number of predicted N-linked glycosylation sites varies from 9 in BOMV to 17 in EBOV and RESTV. Two of these glycosylation sites are situated in the GP2 subunit (N563 and N618) and they are conserved in all ebolavirus species, MARV and LLOV. All remaining sites are located within the GP1 subunit, especially the glycan cap and MLD. Only 4 sites in GP1 are fully conserved in all ebolavirus species: N40 in the base (also present in LLOV), N204 in a flexible loop between the base and head domains, and N257 and N268 in the glycan cap. All other N-linked glycosylation sites in the glycan cap are shared between a smaller set of ebolavirus species, but virtually all N-linked sites within the MLD are unique, in line with the disordered nature and high overall sequence variability of this region. There are 3 predicted C-mannosylation sites conserved in all ebolavirus species and LLOV, but conspicuously missing in MARV. The first WXXW motif is situated in the glycan cap, at W288 (in EBOV), close to the junction with the MLD. In addition, a tandem WXXWXXW motif is situated at W645/W648 in the membrane proximal region of the GP2 subunit.

To visualize the N-glycan shield, we built a pseudomodel of EBOV GP with the core pentasaccharide of each site linked to the corresponding residue of GP1/GP2 (see Figure 1B). The GP trimer forms a chalice-shaped structure with GP2 as the stem, and GP1 as the bowl on top. The conserved sites N40, N204, N257, N268, N563 and N618 are distributed evenly across the structure, whereas the remaining sites are situated primarily at the rim of the bowl extending outwards from the glycan cap. The glycans occupy much of the available surface of GP. Moreover, the disordered MLD connects the tip of the glycan cap with the lower base of the cup and can be expected to further shield the surface of GP. This pseudo model includes only the common core pentasaccharide of the N-linked glycans and it is not known how the glycans are processed in the context of folded GP, as predicted sites are not always glycosylated and the processing from high-mannose precursors to hybrid and mature complex glycans may depend on many unpredictable factors, including local structural constraints. We investigated the patterns of site-specific glycosylation of ebolavirus GP with LC-MS/MS based glycoproteomics experiments, using recombinant soluble ectodomains (GPΔTM) of EBOV and BDBV, as well as the corresponding full-length GP from virus-like particles produced by co-expression with VP40. We compared GPΔTM from human HEK293 and insect S2 cells, both commonly used for structural biology studies, experimental immunizations, antibody selection and serological tests.

Our results cover 14/17 and 17/17 predicted sites in EBOV GP from HEK293 and S2 cells, respectively, as well as 12/14 predicted sites of BDBV GP from both expression platforms (see Figure 2 and Supplementary Table S1). As expected, the N-linked glycosylation patterns of GP from HEK293 and S2 cells are dominated by complex and paucimannose/hybrid glycans, respectively. The glycosylation of GP from especially HEK293 cells is extremely heterogeneous, with some sites carrying over 40 unique glycan compositions. Predicted sites N278 and N391 in BDBV were only detected as unglycosylated asparagines. Most detected glycan compositions are compatible with di-, tri- and tetra-antennary, galactosylated complex glycans with or without a single (core) fucose residue and a variable number of terminal sialic acids, as previously described in glycomics analyses [30, 35]. While complex glycans dominate the overall picture, selected sites show clear and robust enrichment of unprocessed glycans, (*i.e*. high-mannose and hybrid structures in the HEK293-derived samples). These include particularly the conserved N257 and N563 sites, in both EBOV and BDBV GP. In good agreement with these observations in the HEK293-derived samples, N257 and N563 are also enriched in unprocessed high-mannose glycans in the S2-derived samples of both EBOV and BDBV GP, indicating that processing of these sites is somehow structurally restricted. Our pseudomodel of EBOV GP indicates that N257 may be partially buried between the head domain and glycan cap (*i.e*. its first asparagine-linked GlcNAc residue), and N563 similarly between the head domain and GP2. Whereas sites N40/N268/N454 in EBOV GP, and N400/N454 in BDBV GP also show elevated levels of unprocessed glycans in selected samples, we refrain from any conclusions on these sites due to a relatively shallow coverage in the underlying mass spec data and the lack of agreement between HEK293/S2 or EBOV/BDBV samples. Nevertheless, the data clearly indicate a lack of processing at the conserved N257 and N563 sites in both tested ebolavirus species and expression platforms. These findings are confirmed in LC-MS/MS experiments of EBOV and BDBV virus-like particles derived from HEK293 cells, where we also detected a large fraction of high-mannose and hybrid glycans at N257/N563 against a background of highly processed complex glycans at the remaining covered sites (see Supplementary Figures S2 and S3). The abundance of unprocessed glycans is most prominent at site N563, where the vast majority of glycans consists of hybrid and high-mannose forms in both full-length GP and GPΔTM. At site N257, the abundance of unprocessed glycans is markedly lower in the full-length EBOV GP from VLPs.

**Figure 2.**
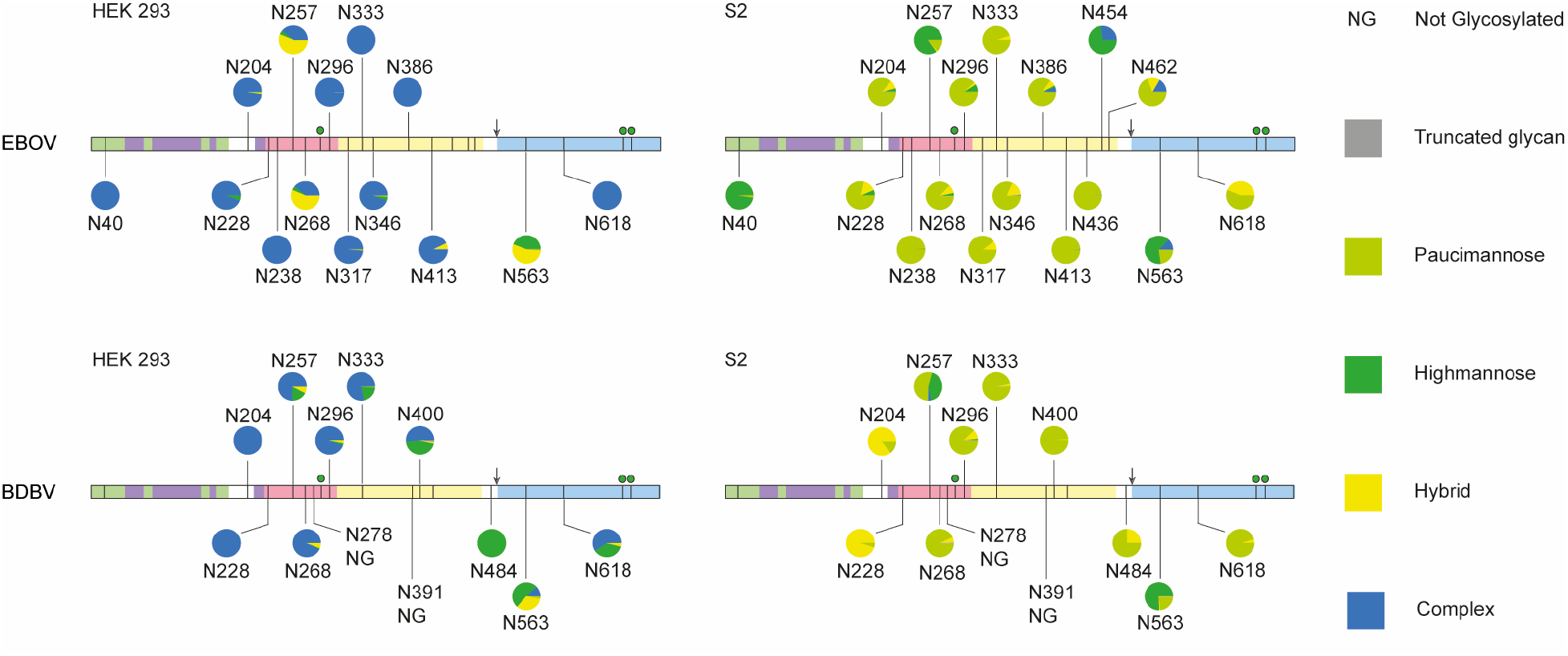
Overview of site-specific N-linked glycan processing in ebolavirus GPΔTM from HEK293 and S2 cells as determined by LC-MS/MS. The glycans were classified by HexNAc content as truncated, paucimannose, high-mannose, hybrid or complex. Shown is the average of a duplicate experiment.

Our experiments also cover the C-mannosylation site at W288 in the glycan cap (see Figure 3). The GPΔTM constructs used here are truncated before the second C-mannosylation motif at W645/W648 and therefore not covered in these experiments. The GP samples derived from HEK293 cells both contain a mixture of C-mannosylated and unmodified W288, with an estimated occupancy of 1-10%. In contrast, this modification is completely absent in both samples derived from S2 cells. The presence of C-mannosylated W288 was confirmed in the glycoproteomics experiments on full-length GP in the virus-like particles formed by co-expression with VP40 (see Supplementary Figure S4). Unfortunately, we could not detect any peptides that cover the second C-mannosylation motif at W645/W648 in these samples.

**Figure 3.**
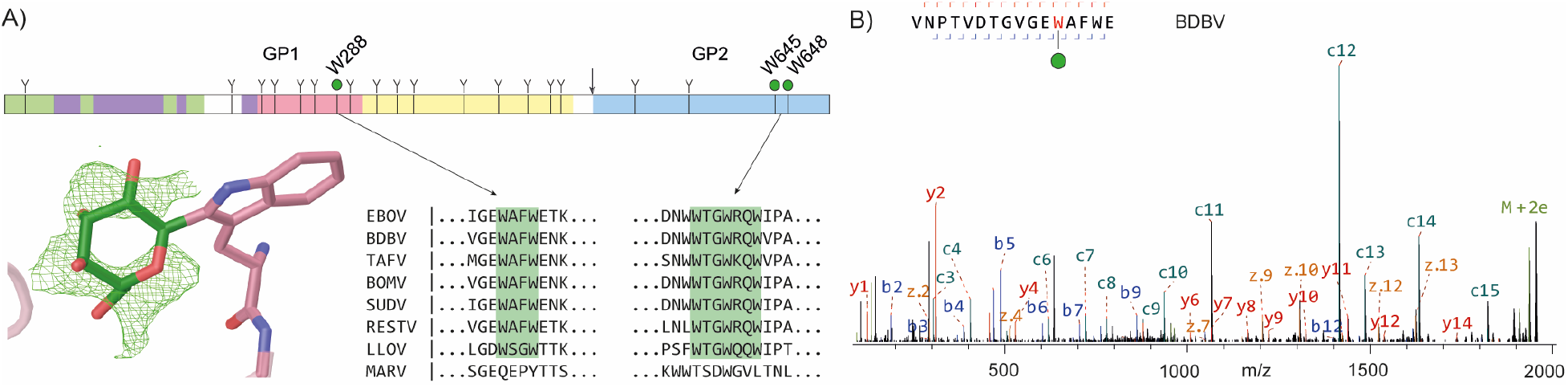
C-mannosylation in ebolavirus GP. A) schematic domain structure ebolavirus GP with highlighted C-mannosylation sites. The modelled structure of C-mannosylated W288 was based on average Fo-Fc density (shown in green) of twelve isomorphic GP crystal structures. B) LC-MS/MS spectrum of C-mannosylated W288 from BDBV GPΔTM. Note the prominent c11-c12 peaks provide direct evidence for the presence and localization of the Hex (+162 Da) modification of W288.

The W288 site is situated in a lesser ordered region of the glycan cap (the β17-β18 loop) just before the start of the MLD, which is deleted in the constructs of most structural studies. While most available GP structures do not model the corresponding region, we identified a set of 12 deposited isomorphic GP crystal structures of HEK293-derived material with electron density for W288 and its adjacent residues [41–45]. The individual crystal structures did not show a clear *F_o_-F_c_* density corresponding to the C2-linked mannose, but after averaging all available electron density maps, a clear ring structure did appear. The weak observed electron density is consistent with the low occupancy of the modification observed in our glycoproteomics experiments. The C2-linked mannose residue was modelled in the extra density, positioning it at the exposed surface of the glycan cap, pointing towards the center of the β17-β18 loop.

We also mapped out the patterns of O-linked glycosylation in ebolavirus GP (see Figure 4). In contrast to N- and C-linked glycosylation, there is no clear sequence motif to predict O-linked glycosylation sites. The modification generally occurs in serine/threonine-rich disordered regions, such as the MLD of filovirus GPs. Whereas the presence of O-glycans in the MLD is well-known, the precise localization of these modifications remains unclear (the MLD contains more than 50 possible S/T residues). For these experiments, we first removed all N-linked glycans by PNGase F digestion. This reduces the complexity of the glycopeptide mixture to facilitate O-linked glycopeptide identification and site localization, while leaving a clear mark at the digested N-glycan site by deamidation of the asparagine residue (resulting in a +1 Da mass shift).

**Figure 4.**
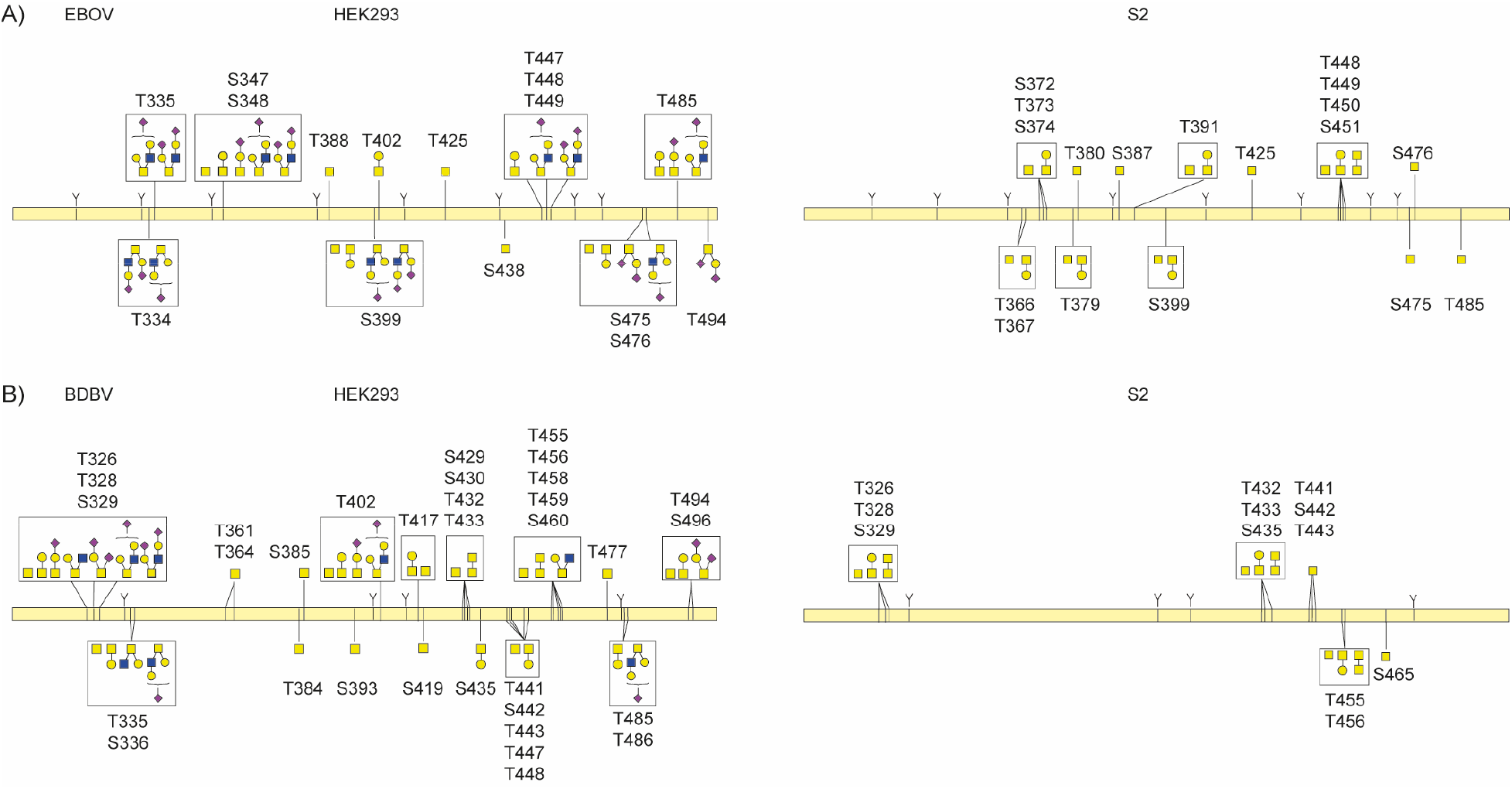
O-linked glycosylation in the MLD of ebolavirus GPΔTM from EBOV (A) and BDBV (B). Glycans drawn within a box represent multiple detected compositions per site. Glycans connected to multiple indicated sites could not be unambiguously localized from the LC-MS/MS data.

In EBOV GP, we detected 12 unique O-linked glycosylation sites in the MLD of GPΔTM from HEK293 cells versus 12 in S2 cells, with 5 sites in common. In BDBV GP we detected 16 unique O-linked glycosylation sites in the MLD of GPΔTM from HEK293 cells versus 5 in S2 cells, with 4 sites in common. Whereas O-linked glycosylation was dominated by simple Tn antigen and core 1 structures in samples from S2 cells, samples from HEK293 cells also contained extended and sialylated core 1 and core 2 structures, especially in EBOV GP. Multiple unique glycan compositions were often detected for a given site, further adding to the extreme heterogeneity of GP due to its glycosylation.

We also detected several O-glycosylation sites outside the MLD of both EBOV and BDBV GP (see Figure 5). In BDBV GP we detected O-linked glycosylation at T280, with 6 unique glycan compositions amounting to an estimated total occupancy of ~10%. This threonine residue is part of a putative NPT glycosylation sequon, but we only detect the unmodified asparagine, which remains unprocessed presumably because of the following proline residue. The modified threonine is shared only by BDBV and TAFV GP, but absent in EBOV, SUDV, BOMV, RESTV, MARV and LLOV. We also detected O-linked glycosylation at T206 in EBOV GP, with 6 unique glycan compositions and an estimated occupancy of ~5%. This threonine is part of the glycosylation sequon of N204, which is fully occupied by N-glycans as evidenced by the deamidated asparagine and the N-linked glycoproteomics data discussed earlier. The N-linked glycosylation sequon including the modified threonine is conserved among all ebolavirus species, but does not exist in MARV and LLOV. Residues adjacent to this sequon show substantial variation between ebolavirus species and modified T206 was not detected in the BDBV GP samples. The close juxtaposition of N- and O-glycans is also observed in the MLD of both EBOV and BDBV GP, where T335 is part of the N-linked glycosylation sequon of N333 and detected in GPΔTM from both species as an O-linked glycosylation site. Similarly, S348/T388/S438 in EBOV GP and T402/T488 in BDBV GP are all part of N-glycosylation sequons. Finally, we also observed O-linked glycosylation within the strep-tag of the constructs (see Supplementary Figure S5). The presence of the O-linked glycans at T206 (EBOV) and T280 (BDBV) outside the MLD could be confirmed in our glycoproteomics measurements of full-length GP from virus-like particles (see Supplementary Figure S4).

**Figure 5.**
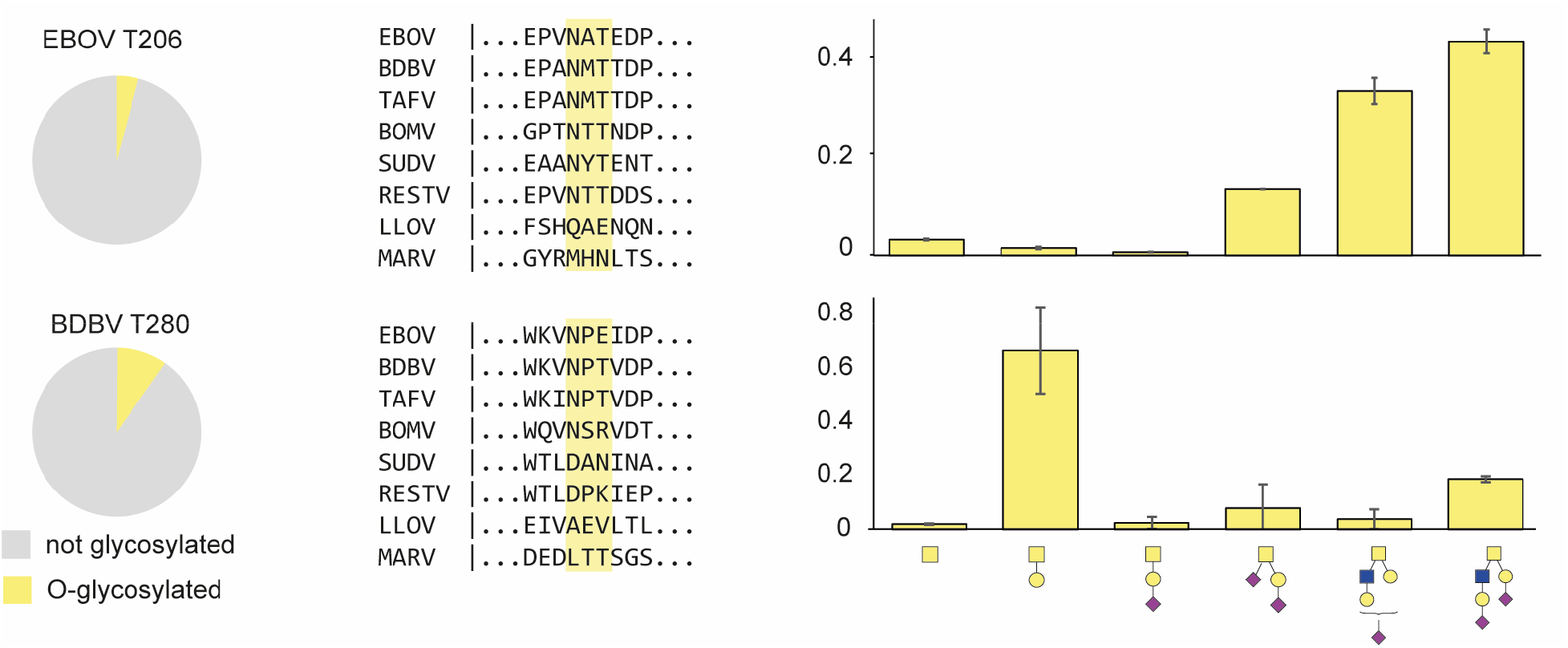
O-linked glycosylation outside the MLD in ebolavirus GP. (left) pie charts represent the occupancy of the O-linked modification. (middle) sequence conservation of the detected O-linked glycosylation sites. (right) distribution of glycan types at the indicated sites (average +/- stdev from duplicate experiments)

This high extent of glycosylation must be accommodated by antibodies against ebolavirus GP. To understand the contribution of glycans to neutralization epitopes (and restrictions they impose) we screened the Protein Data Bank for structural models of ebolavirus GP in complex with neutralizing antibodies and also looked for linear B-cell epitopes reported in literature that span the glycosylation sites we detected in our experiments (see Figure 6A) [46–56]. The glycan-rich epitopes we report in this overview include both cases of direct contacts between modelled GP glycans and CDR residues, as well as brushing interactions of adjacent glycans with the framework regions of the variable domains, which may sterically restrict binding. It should be noted that glycans are typically incompletely modelled in GP-antibody structures and that inference of these brushing interactions is not an exact determination. This overview indicates that neutralizing antibodies span a broad range of epitopes that cover essentially all N-linked glycosylation sites (N204 and the entire MLD have not yet been modelled in structural studies and are therefore not represented in this analysis). The conserved GP1 glycans (N40, N257, and N268) all contribute to the epitopes of neutralizing antibodies, with possible glycan-antibody interactions for N268 reported in the epitopes of as many as 7 unique monoclonal antibodies. This includes the therapeutic monoclonal antibody Mab114, which makes additional contacts with N238. The components of the therapeutic ZMapp mixture (c2G4, c4G7 and c13C6) also interact with N40, N238, N268 and N563. The epitopes of the three components in the therapeutic REGN-EB3 mixture are not defined to atomic detail (and therefore not included in the overview), but published negative stain EM reconstructions suggest possible interaction with N563 and the glycan cap [57].

**Figure 6.**
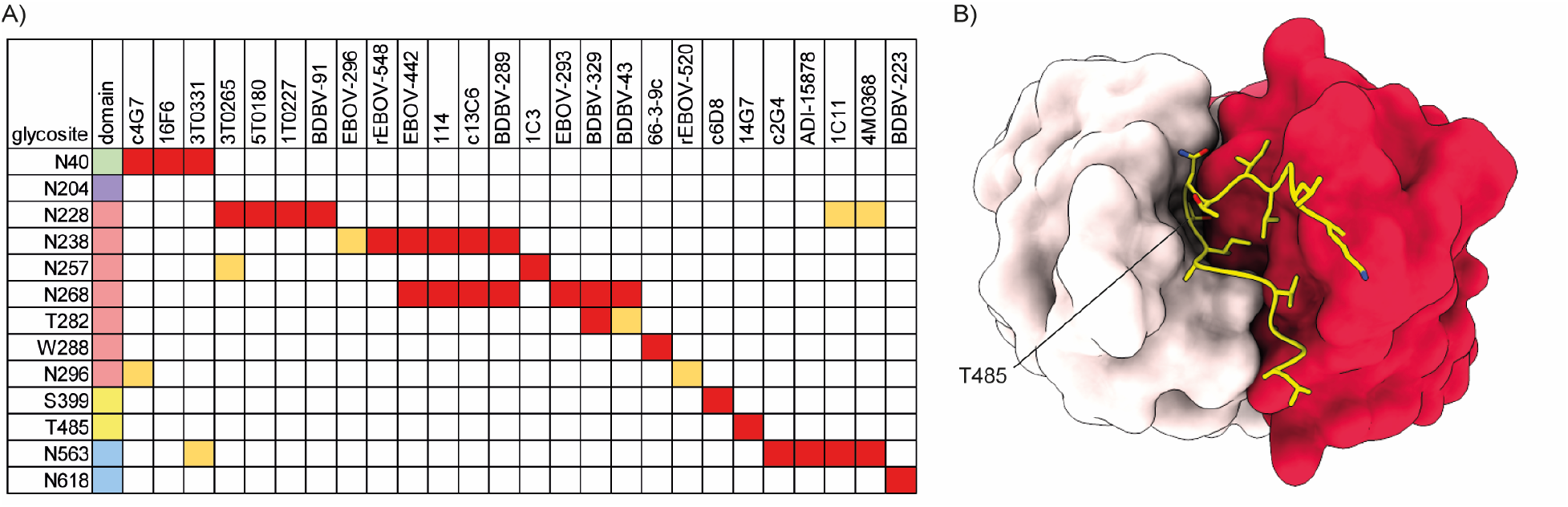
Glycan-containing epitopes of monoclonal antibodies against ebolavirus GP. A) overview of glycosylation sites associated with indicated epitopes. Direct contacts and clashes with CDRs are indicated in red, brushing interactions with framework regions in light orange. B) highlighted structure of 14G7 in complex with a linear epitope from the MLD (PDB ID:2Y6S), showing that the O-linked glycosylation site T485 is deeply buried in the cleft between heavy (dark red) and light chain (light pink).

Besides N-glycans, we also noted several putative interactions with C-mannosylated W288 and O-linked glycosylation sites. The monoclonal antibodies BDBV-329 and BDBV-43 are in close proximity to T280 with their CDRH3 and framework 3 regions, respectively. The monoclonal antibody 66-3-9c binds a linear epitope that spans the C-mannosylation site W288. Similarly, c6D8 binds a linear epitope that spans the O-linked glycosylation site S399. The monoclonal antibody 14G7 binds to a linear epitope in the MLD that spans the O-linked glycosylation site T485. A crystal structure of the 14G7 Fab in complex with its unglycosylated epitope reveals that T485 is buried deep within the cleft between heavy and light chains, where it is in direct contact with CDRH3 residues (see Figure 6B). It is therefore unlikely to accommodate the bulky O-glycans detected in our experiments, further highlighting the potential epitope shielding effects of not just N-glycans, but also O-glycans in the MLD.

## Discussion

Here we have presented a comprehensive overview of glycosylation in ebolavirus GP, using glycoproteomics to resolve the patterns of site-specific N-, O- and C-linked glycans. In the GP samples derived from HEK293 cells, we observed heterogeneous, complex N-glycosylation overall, but noted enrichment of unprocessed glycans at two conserved sites N257 and N563. It is known that ebolavirus GP interacts with cell surface lectins DC-SIGN/L-SIGN in a high-mannose glycan dependent manner [27–29]. Our results indicate that the unprocessed glycans present at N257 and N563 may be primarily responsible for the interaction, thereby facilitating host cell attachment and infectivity.

We also detected up to 16 unique site-specific O-glycans in the MLD of GP, revealing a heterogeneous mixture of not only simple Tn-antigen (*i.e*. a single GalNAc), but also extended sialylated core 1 and core 2 structures. The sites detected in our experiments are likely just the tip of the iceberg, as the dense decoration of the MLD with O-glycans may make proteolytic digestion of the MLD especially difficult and the presence of multiple glycans in the same peptide makes it exponentially more challenging to confidently make assignments from the raw LC-MS/MS data.

We further confirmed the presence of the C-mannosylation site in GP at W288, which is completely conserved in all ebolavirus species and LLOV, but not MARV. The second motif at W645/W658 is also missing in MARV, but whereas the MARV sequence at W288 completely diverges from the ebolavirus species, the MARV region corresponding to W645/W648 is similarly rich in tryptophan residues (see Figure 3A). The MARV GP sequence in this region is thereby primed to acquire a C-mannosylation motif through a single deletion or a tryptophan substitution at any of four adjacent positions. Conversely, this could also indicate that the C-mannosylation motifs were present in a common ancestor with ebolaviruses and LLOV but lost in MARV. Whereas C-mannosylation is known to be important for the stability and folding of secreted human glycoproteins [38, 39], its role in ebolavirus replication and pathogenesis remains unclear.

The presence of heterogeneous N-, O- and C-linked glycosylation add up to a staggering complexity of GP composition. In the case of EBOV GP, the 17 N-glycosylation sites alone, each linked to on average a dozen unique glycan compositions, already give rise to an enormous number of permutations. Add to this the heterogeneity of O-linked glycosylation and a picture emerges where no two copies of GP on a virion are strictly identical. Meanwhile, the glycans represent a major component of the overall GP structure. The 17 N-glycans already contribute approximately one quarter of the molecular weight of GP, all situated at its exposed surface (counting ~74 kDa of polypeptide and on average 1.5 kDa per N-glycan). Although antibodies evidently mount a neutralizing response to infection or vaccination, the high variability of the exposed GP surface due to heterogeneous glycosylation must frustrate the overall binding efficiency of antibodies that accommodate glycans in their epitope and restrict good binders to only the core elements of glycans that are common between the countless variations of GP present on the surface of mature virions. From this perspective the glycans contribute to immune evasion not only by sterically shielding neutralization epitopes, but also by blurring the molecular identity of the envelope glycoprotein.

Glycoproteomics studies on other envelope glycoproteins from divergent virus species, such as HIV-1, Lassa virus, MERS-CoV, SARS-CoV-2 and the herpesviruses, all show a similar trend of heterogeneous complex glycosylation with unprocessed glycans at selected sites [58–64]. From studies on HIV-1 gp120 it has been shown that the lack of processing of certain N-glycans is caused by local crowding and reduced accessibility to processing enzymes, resulting in enrichment of high-mannose glycans [65–68]. Neither glycan at N257 and N563 in ebolavirus GP fits this description, but both have their first N-linked GlcNAc residue partially buried in interactions with surrounding side chains. An intriguing possibility is that these interactions at the base of the glycan limit its conformational degrees of freedom to negatively impact processing at the antennae. Whereas N-linked glycosylation is universally known to play a role in the replication cycles of enveloped viruses, O-linked glycosylation is less well-studied and perhaps less common. Recent studies on a range of herpesvirus glycoproteins, SARS-CoV-2 Spike, the attachment proteins G of paramyxoviruses, and hepatitis C virus E2 point to a role in envelope glycoprotein processing, trafficking, host cell attachment, and immune evasion [58, 59, 69–72].

All three types of glycosylation observed in ebolavirus GP will alter its antigenic surface. The overview provided in Figure 6 illustrates how the site-specific glycosylation observed in our experiments contributes to and restricts the epitopes of currently known neutralizing antibodies. Whereas bulkier types of glycans at specific sites may indeed modulate the binding affinity of the indicated antibodies, the presented overview shows quite the opposite of the glycans’ shielding effects in ebolavirus GP. That interpretation would be a kind of survivorship bias, as the monoclonal antibodies have been selected for binding and neutralization. The effective shielding would perhaps be better illustrated by the antibodies that don’t bind or neutralize ebolavirus infection because of the steric clashes with glycans. Several studies have illustrated this by showing greatly enhanced sensitivity of (pseudo) ebolavirus to serum neutralization after glycan removal by mutagenesis [31, 32].

Future studies may shed light on how the heterogenous glycan composition of GP may modulate antibody binding. Similarly, the close juxtaposition of N- and O-linked glycan raises the intriguing possibility of an interplay between the two types of modifications. Furthermore, the exact role of C-mannosylation and O-linked glycans outside the MLD also remain to be investigated. Nevertheless, we have presented a comprehensive overview of ebolavirus GP glycosylation that may provide a useful framework for future immunological and structural studies on GP-antibody interactions.

## Acknowledgements

The authors would like to thank everyone in the Biomolecular Mass Spectrometry and Proteomics group at Utrecht University for support and helpful discussions. This research was funded by the Dutch Research Council NWO Gravitation 2013 BOO, Institute for Chemical Immunology (ICI; 024.002.009) to JS, and NIAID U19 AI142790 to EOS.

## Materials and Methods

### Ebolavirus GP sequence analysis

The indicated full-length GP reference sequences were downloaded from UniProt. The LLOV-GP sequence had to be reconstructed from the two separate GP1 and GP2 entries in UniProt. Sequence alignment was performed with ClustalX 2.1 [73]. The sequence IDs and resulting alignment are provided in the Supplementary Information. The cladogram was generated with FigTree (version 1.4.4). N-linked glycosylation sites were predicted by identifying all NXS/T sequences with NetNGlyc-1.0. C-mannosylation sites were predicted by identifying all WXXW sequences by manual inspection.

### Pseudomodel building of glycosylated EBOV GP

A homology model EBOV GP (strain Mayinga ‘76) was generated with SWISS-MODEL to fill in missing loops using PDB ID 5jq3 as a template [45, 74]. The core pentasaccharides were added to GP1 and GP2 subunits separately with GLYCAM Glycoprotein Builder (GLYCAM Web, Woods group 2021). The full trimer was reconstructed by alignment with the biological assembly of 5jq3. The loop containing N204 was manually removed because it produced clashes with neighboring subunits in the full trimer. The figures were generated with ChimeraX 1.2.5 [75].

### Glycan-containing epitopes of monoclonal antibodies

Structures of monoclonal antibodies in complex with ebola virus GP were retrieved from the PDB (with PDB IDs: 2y6s, 3s88, 5fhc, 5kel, 5kem, 5ken, 6ea7, 6n7j, 6pci, 6qd7, 6qd8, 6s8d, 7kej, 7kew, 7kex, 7kfe, 7kf9 and 7kfb) [46–56]. The structures were aligned with the glycosylated ZEBOV GP pseudomodel described above, using the MatchMaker function of ChimeraX 1.2.5, and glycans within 6 Å of the CDRs or framework regions of the modelled antibodies were included in the overview.

### Modelling of C-mannosylated W288

identified 12 isomorphous published crystal structures of HEK293-derived EBOV GP samples in the PDB (PDB codes 6f6n, 6f6i, 6f54, 6nae, 5jqb, 5jq7, 5jq3, 6g9b, 6g9i, 6g95, 6hro and 6hs4) [41–45]. To obtain higher signal-to-noise ratios from these maps, the 2*F*_o_-*F*_c_ and *F*_o_-*F*_c_ difference maps were averaged using COOT [76]. This averaged map showed clear ring-shaped electron density next to W288 in the *F*_o_-*F*_c_ difference map at a contour level of 2.7 root mean square deviation. A C-mannosyl group was modelled next to W288 using the EBOV GP structure 6hs4 as a template in COOT. To accommodate realistic geometry, the tryptophan had to be repositioned slightly, albeit still in agreement with the local electron density. Care was taken to model the mannose with a ring-flipped ^1^C_4_ chair conformation [77–79].

### Production and purification of ebola virus GP ectodomains

EBOV and BDBV GP were produced by both transient transfection of HEK293T cells and stable transfection of Drosophila melanogaster S2 cells. Lipofectamine 3000 (Invitrogen) was used to transfect HEK293T cells, and Effectene (Qiagen) was used to produce stable S2 cells with a modified pMT-puro vector plasmid containing the GP gene of interest and stable selection of transfected cells with 6 μg/mL puromycin. HEK293T cells were grown at 37°C with 5% CO2 in DMEM media (Gibco) supplemented with 10% FBS in T75 flasks and expanded into 10-stack flasks (Corning) for transfection. S2 cells were selected at 27°C in complete Schneider’s medium and then transferred to Insect Xpress medium (Lonza) for large-scale expression in 2-liter Erlenmeyer flasks. Secreted GP ectodomain expression was induced with 500 mM CuSO4, and supernatant harvested after 4 days. Ebola virus GP was engineered with a double Strep-tag at the C terminus to facilitate purification using Strep-trap HP 5mL column (GE) and then further purified by Superdex 200 size exclusion chromatography (SEC) in 25 mM Tris-buffered saline (Tris-HCl, pH 7.5, 150 mM NaCl [TBS]).

### Production and purification of ebolavirus-like particles

EBOV and BDBV virus-like particles were produced by transfecting HEK293T cells. Polyethylenimine (PEI) was used to transfect HEK293T cells with a modified phCMV plasmid containing the full-length GP gene of interest and a modified pTriEx plasmid containing the full-length EBOV VP40 gene at a 2:5 ratio (w:w), respectively. The VLP supernatant was clarified by centrifugation after 48 hours. The clarified supernatant was further purified using a 20% sucrose cushion ultra-centrifuge spin at 106,800xg for 3hrs. The cushion and supernatant was carefully decanted and the pellet washed with sterile PBS 2 times. Following the wash, the pellet was incubated overnight in 0.75mL of PBS and resuspended.

### Glycoproteomics sample preparation

For N-linked glycan analysis, the recombinant GP was denatured at 95 °C in a final concentration of 2% sodium deoxycholate (SDC), 200 mM Tris/HCl, 10 mM tris(2-carboxyethyl)phosphine, pH 8.0 for 10 min followed with 30 min reduction at 37 °C for 30 min. Samples were next alkylated by adding 40 mM iodoacetamide and incubated in the dark at room temperature for 45 min. 3 μg recombinant GP was used for each protease digestion. Samples were split in three for parallel digestion with trypsin (Promega), alpha lytic protease (Sigma), and gluC (Sigma)-trypsin. For each protease digestion, 18 μL of the denatured, reduced, and alkylated samples was diluted in a total volume of 100 μL 50 mM ammonium bicarbonate, adding proteases in a 1:15 ratio (w:w) for incubation overnight at 37 °C. For the gluC-trypsin digestion, gluC was added first for two hours, followed by incubation with trypsin overnight. After overnight digestion SDC was removed through precipitation by adding 2 μL formic acid (FA) and centrifugation at 14,000 rpm for 20 min. Following centrifugation, the supernatant containing the peptides was collected for desalting on a 30 μm Oasis HLB 96-well plate (Waters). The Oasis HLB sorbent was activated with 100% acetonitrile and subsequently equilibrated with 10% formic acid in water. Next, peptides were bound to the sorbent, washed twice with 10% formic acid in water and eluted with 100 μL of 50% acetonitrile/5% formic acid in water (v/v). The eluted peptides were vacuum-dried and resuspended in 100 μL of 2% formic acid in water. For O-linked glycan analysis, the recombinant GP was first treated with PNGase F (Sigma) to remove N-glycans. 4 μL PNGase F was added to the sample in PBS and incubated at 37 °C overnight. Following N-glycan removal, GPs were digested following the same protocol as for N-linked glycan analysis, using parallel digestion with trypsin and aLP. Both N- and O-linked analyses were performed in duplicate.

### Glycoproteomics LC-MS/MS measurements

For each sample and protease digestion, approximately 0.15 μg of peptides were run by online reversed phase chromatography on an Agilent 1290 UHPLC or Dionex UltiMate 3000 (Thermo Fisher Scientific) coupled to a Thermo Scientific Orbitrap Fusion mass spectrometer. A Poroshell 120 EC C18 (50 cm x 75 μm, 2.7 μm, Agilent Technologies) analytical column and a ReproSil-Pur C18 (2 cm x 100 μm, 3 μm, Dr. Maisch) trap column were used for peptide separation. The duplicate samples were analyzed with two different mass spectrometry methods, using identical LC-MS parameters and distinct fragmentation schemes. In one method, peptides were subjected to Electron Transfer/Higher-Energy Collision Dissociation fragmentation. In the other method, all precursors were subjected to HCD fragmentation, with additional EThcD fragmentation triggered by the presence of glycan reporter oxonium ions. A 90-min LC gradient from 0% to 44% acetonitrile was used to separate peptides at a flow rate of 300 nl/min. Data was acquired in data-dependent mode. Orbitrap Fusion parameters for the full scan MS spectra were as follows: a standard AGC target at 60 000 resolution, scan range 350-2000 m/z, Orbitrap maximum injection time 50 ms. The ten most intense ions (2+ to 8+ ions) were subjected to fragmentation. For the EThcD fragmentation scheme, the supplemental higher energy collision dissociation energy was set at 27%. MS2 spectra were acquired at a resolution of 30,000 with an AGC target of 800%, maximum injection time 250 ms, scan range 120-4000 m/z and dynamic exclusion of 16 s. For the triggered HCD-EThcD method, the LC gradient and MS1 scan parameters were identical. The ten most intense ions (2+ to 8+) were subjected to HCD fragmentation with 30% normalized collision energy from 120-4000 m/z at 30,000 resolution with an AGC target of 100% and a dynamic exclusion window of 16 s. Scans containing any of the following oxonium ions within 20 ppm were followed up with additional EThcD fragmentation with 27% supplemental HCD fragmentation. The triggering reporter ions were: Hex(1) (129.039; 145.0495; 163.0601), PHex(1) (243.0264; 405.0793), HexNAc(1) (138.055; 168.0655; 186.0761), Neu5Ac(1) (274.0921; 292.1027), Hex(1) HexNAc(1) (366.1395), HexNAc(2) (407.166), dHex(1)Hex(1)HexNAc(1) (512.1974), and Hex(1)HexNAc(1)Neu5Ac(1) (657.2349). EThcD spectra were acquired at a resolution of 30,000 with a normalized AGC target of 400%, maximum injection time 250 ms, and scan range 120-4000 m/z.

### Glycoproteomics data analysis

The acquired data was analysed using Byonic (v3.9.6 [80]) against a custom database of recombinant ebola virus GP protein sequences and the proteases used in the experiment, searching for glycan modifications with 12/24 ppm search windows for MS1/MS2, respectively. Up to six missed cleavages were permitted using C-terminal cleavage at R/K for trypsin, R/K/E/D for gluC-trypsin, or T/A/S/V for alpha lytic protease. For N-linked analysis, carbamidomethylation of cysteine was set as fixed modification, oxidation of methionine/tryptophan as variable common 1, and hexose on tryptophan as variable rare 1. N-glycan modifications were set as variable common 2, allowing up to max. 2 variable common and 1 rare modification per peptide. All N-linked glycan databases from Byonic were merged into a single non-redundant list to be included in the database search. All reported glycopeptides in the Byonic result files were manually inspected for quality of fragment assignments. All glycopeptide identifications were merged into a single non-redundant list per sequon. Glycans were classified based on HexNAc content as truncated (≤ 2 HexNAc; < 3 Hex), paucimannose (2 HexNAc, 3 Hex), high-mannose (2 HexNAc; > 3 Hex), hybrid (3 HexNAc) or complex (> 3 HexNAc). Byonic search results were exported to mzIdentML format to build a spectral library in Skyline (v20.1.0.31 [81]) and extract peak areas for individual glycoforms from MS1 scans. The full database of variable N-linked glycan modifications from Byonic was manually added to the Skyline project file in XML format. Reported peak areas were pooled based on the number of HexNAc, Fuc or NeuAc residues to distinguish truncated, paucimannose, high-mannose, hybrid, and complex glycosylation, or the degree of fucosylation and sialylation, respectively. For O-linked analysis, all the same protease digestion parameters and peptide modifications were used, with the addition of deamidation at asparagine/glutamine as variable rare 1. O-glycan modifications were set as variable common 6, allowing a maximum of 6 variable common and 2 rare modifications per peptide.

### Data Availability

The raw LC-MS/MS files and glycopeptide identifications have been deposited to the ProteomeXchange Consortium via the PRIDE partner repository with the dataset identifier PXD031459. All reagents and relevant data are available from the authors upon request.

## References

1. Burk, R., L. Bollinger, J.C. Johnson, J. Wada, S.R. Radoshitzky, G. Palacios, S. Bavari, P.B. Jahrling and J.H. Kuhn, Neglected filoviruses. FEMS microbiology reviews, 2016. 40(4): p. 494–519.

2. Jacob, S.T., I. Crozier, W.A. Fischer, A. Hewlett, C.S. Kraft, M.-A. de La Vega, M.J. Soka, V. Wahl, A. Griffiths and L. Bollinger, Ebola virus disease. Nature reviews Disease primers, 2020. 6(1): p. 1–31.

3. Leroy, E.M., B. Kumulungui, X. Pourrut, P. Rouquet, A. Hassanin, P. Yaba, A. Délicat, J.T. Paweska, J.-P. Gonzalez and R. Swanepoel, Fruit bats as reservoirs of Ebola virus. Nature, 2005. 438(7068): p. 575–576.

4. Marí Saéz, A., S. Weiss, K. Nowak, V. Lapeyre, F. Zimmermann, A. Düx, H.S. Kühl, M. Kaba, S. Regnaut and K. Merkel, Investigating the zoonotic origin of the West African Ebola epidemic. EMBO molecular medicine, 2015. 7(1): p. 17–23.

5. Goldstein, T., S.J. Anthony, A. Gbakima, B.H. Bird, J. Bangura, A. Tremeau-Bravard, M.N. Belaganahalli, H.L. Wells, J.K. Dhanota and E. Liang, The discovery of Bombali virus adds further support for bats as hosts of ebolaviruses. Nature microbiology, 2018. 3(10): p. 1084–1089.

6. Amman, B.R., S.A. Carroll, Z.D. Reed, T.K. Sealy, S. Balinandi, R. Swanepoel, A. Kemp, B.R. Erickson, J.A. Comer and S. Campbell, Seasonal pulses of Marburg virus circulation in juvenile Rousettus aegyptiacus bats coincide with periods of increased risk of human infection. 2012.

7. Swanepoel, R., S.B. Smit, P.E. Rollin, P. Formenty, P.A. Leman, A. Kemp, F.J. Burt, A.A. Grobbelaar, J. Croft and D.G. Bausch, Studies of reservoir hosts for Marburg virus. Emerging infectious diseases, 2007. 13(12): p. 1847.

8. Towner, J.S., X. Pourrut, C.G. Albariño, C.N. Nkogue, B.H. Bird, G. Grard, T.G. Ksiazek, J.-P. Gonzalez, S.T. Nichol and E.M. Leroy, Marburg virus infection detected in a common African bat. PloS one, 2007. 2(8): p. e764.

9. Team, W.E.R., Ebola virus disease in West Africa—the first 9 months of the epidemic and forward projections. New England Journal of Medicine, 2014. 371(16): p. 1481–1495.

10. Hoenen, T., A. Groseth and H. Feldmann, Therapeutic strategies to target the Ebola virus life cycle. Nature Reviews Microbiology, 2019. 17(10): p. 593–606.

11. Lee, J.E. and E.O. Saphire, Ebolavirus glycoprotein structure and mechanism of entry. Future virology, 2009. 4(6): p. 621–635.

12. Mehedi, M., D. Falzarano, J. Seebach, X. Hu, M.S. Carpenter, H.-J. Schnittler and H. Feldmann, A new Ebola virus nonstructural glycoprotein expressed through RNA editing. Journal of virology, 2011. 85(11): p. 5406–5414.

13. Volchkova, V.A., H. Feldmann, H.-D. Klenk and V.E. Volchkov, The nonstructural small glycoprotein sGP of Ebola virus is secreted as an antiparallel-orientated homodimer. Virology, 1998. 250(2): p. 408–414.

14. Aleksandrowicz, P., A. Marzi, N. Biedenkopf, N. Beimforde, S. Becker, T. Hoenen, H. Feldmann and H.-J. Schnittler, Ebola virus enters host cells by macropinocytosis and clathrin-mediated endocytosis. The Journal of infectious diseases, 2011. 204(suppl_3): p. S957–S967.

15. Mulherkar, N., M. Raaben, J.C. de la Torre, S.P. Whelan and K. Chandran, The Ebola virus glycoprotein mediates entry via a non-classical dynamin-dependent macropinocytic pathway. Virology, 2011. 419(2): p. 72–83.

16. Nanbo, A., M. Imai, S. Watanabe, T. Noda, K. Takahashi, G. Neumann, P. Halfmann and Y. Kawaoka, Ebolavirus is internalized into host cells via macropinocytosis in a viral glycoprotein-dependent manner. PLoS pathogens, 2010. 6(9): p. e1001121.

17. Saeed, M.F., A.A. Kolokoltsov, T. Albrecht and R.A. Davey, Cellular entry of ebola virus involves uptake by a macropinocytosis-like mechanism and subsequent trafficking through early and late endosomes. PLoS pathogens, 2010. 6(9): p. e1001110.

18. Gong, X., H. Qian, X. Zhou, J. Wu, T. Wan, P. Cao, W. Huang, X. Zhao, X. Wang and P. Wang, Structural insights into the Niemann-Pick C1 (NPC1)-mediated cholesterol transfer and Ebola infection. Cell, 2016. 165(6): p. 1467–1478.

19. Miller, E.H., G. Obernosterer, M. Raaben, A.S. Herbert, M.S. Deffieu, A. Krishnan, E. Ndungo, R.G. Sandesara, J.E. Carette and A.I. Kuehne, Ebola virus entry requires the host-programmed recognition of an intracellular receptor. The EMBO journal, 2012. 31(8): p. 1947–1960.

20. Wang, H., Y. Shi, J. Song, J. Qi, G. Lu, J. Yan and G.F. Gao, Ebola viral glycoprotein bound to its endosomal receptor Niemann-Pick C1. Cell, 2016. 164(1-2): p. 258–268.

21. Saphire, E.O., S.L. Schendel, B.M. Gunn, J.C. Milligan and G. Alter, Antibody-mediated protection against Ebola virus. Nature immunology, 2018. 19(11): p. 1169–1178.

22. Volchkov, V.E., H. Feldmann, V.A. Volchkova and H.-D. Klenk, Processing of the Ebola virus glycoprotein by the proprotein convertase furin. Proceedings of the National Academy of Sciences, 1998. 95(10): p. 5762–5767.

23. Sanchez, A., Z.-Y. Yang, L. Xu, G.J. Nabel, T. Crews and C.J. Peters, Biochemical analysis of the secreted and virion glycoproteins of Ebola virus. Journal of virology, 1998. 72(8): p. 6442–6447.

24. Ito, H., S. Watanabe, A. Sanchez, M.A. Whitt and Y. Kawaoka, Mutational analysis of the putative fusion domain of Ebola virus glycoprotein. Journal of virology, 1999. 73(10): p. 8907–8912.

25. Malashkevich, V.N., B.J. Schneider, M.L. McNally, M.A. Milhollen, J.X. Pang and P.S. Kim, Core structure of the envelope glycoprotein GP2 from Ebola virus at 1.9-Å resolution. Proceedings of the National Academy of Sciences, 1999. 96(6): p. 2662–2667.

26. Weissenhorn, W., A. Carfí, K.-H. Lee, J.J. Skehel and D.C. Wiley, Crystal structure of the Ebola virus membrane fusion subunit, GP2, from the envelope glycoprotein ectodomain. Molecular cell, 1998. 2(5): p. 605–616.

27. Alvarez, C.P., F. Lasala, J. Carrillo, O. Muñiz, A.L. Corbí and R. Delgado, C-type lectins DC-SIGN and L-SIGN mediate cellular entry by Ebola virus in cis and in trans. Journal of virology, 2002. 76(13): p. 6841–6844.

28. Lin, G., G. Simmons, S. Pöhlmann, F. Baribaud, H. Ni, G.J. Leslie, B.S. Haggarty, P. Bates, D. Weissman and J.A. Hoxie, Differential N-linked glycosylation of human immunodeficiency virus and Ebola virus envelope glycoproteins modulates interactions with DC-SIGN and DC-SIGNR. Journal of virology, 2003. 77(2): p. 1337–1346.

29. Simmons, G., J.D. Reeves, C.C. Grogan, L.H. Vandenberghe, F. Baribaud, J.C. Whitbeck, E. Burke, M.J. Buchmeier, E.J. Soilleux and J.L. Riley, DC-SIGN and DC-SIGNR bind ebola glycoproteins and enhance infection of macrophages and endothelial cells. Virology, 2003. 305(1): p. 115–123.

30. Collar, A.L., E.C. Clarke, E. Anaya, D. Merrill, S. Yarborough, S.M. Anthony, J.H. Kuhn, C. Merle, M. Theisen and S.B. Bradfute, Comparison of N-and O-linked glycosylation patterns of ebolavirus glycoproteins. Virology, 2017. 502: p. 39–47.

31. Dowling, W., E. Thompson, C. Badger, J.L. Mellquist, A.R. Garrison, J.M. Smith, J. Paragas, R.J. Hogan and C. Schmaljohn, Influences of glycosylation on antigenicity, immunogenicity, and protective efficacy of ebola virus GP DNA vaccines. Journal of virology, 2007. 81(4): p. 1821–1837.

32. Iraqi, M., A. Edri, Y. Greenshpan, K. Kundu, P. Bolel, A. Cahana, A. Ottolenghi, R. Gazit, L. Lobel and A. Braiman, N-Glycans mediate the ebola virus-GP1 shielding of ligands to immune receptors and immune evasion. Frontiers in cellular and infection microbiology, 2020. 10: p. 48.

33. Jeffers, S.A., D.A. Sanders and A. Sanchez, Covalent modifications of the Ebola virus glycoprotein. Journal of virology, 2002. 76(24): p. 12463–12472.

34. Lennemann, N.J., B.A. Rhein, E. Ndungo, K. Chandran, X. Qiu and W. Maury, Comprehensive functional analysis of N-linked glycans on Ebola virus GP1. MBio, 2014. 5(1): p. e00862–13.

35. Ritchie, G., D.J. Harvey, U. Stroeher, F. Feldmann, H. Feldmann, V. Wahl-Jensen, L. Royle, R.A. Dwek and P.M. Rudd, Identification of N-glycans from Ebola virus glycoproteins by matrix-assisted laser desorption/ionisation time-of-flight and negative ion electrospray tandem mass spectrometry. Rapid Communications in Mass Spectrometry: An International Journal Devoted to the Rapid Dissemination of Up-to-the-Minute Research in Mass Spectrometry, 2010. 24(5): p. 571–585.

36. Wang, B., Y. Wang, D.A. Frabutt, X. Zhang, X. Yao, D. Hu, Z. Zhang, C. Liu, S. Zheng and S.-H. Xiang, Mechanistic understanding of N-glycosylation in Ebola virus glycoprotein maturation and function. Journal of Biological Chemistry, 2017. 292(14): p. 5860–5870.

37. Lennemann, N.J., M. Walkner, A.R. Berkebile, N. Patel and W. Maury, The role of conserved N-linked glycans on Ebola virus glycoprotein 2. The Journal of infectious diseases, 2015. 212(suppl_2): p. S204–S209.

38. Furmanek, A. and J. Hofsteenge, Protein C-mannosylation: facts and questions. Acta biochimica polonica, 2000. 47(3): p. 781–789.

39. Julenius, K., NetCGlyc 1.0: prediction of mammalian C-mannosylation sites. Glycobiology, 2007. 17(8): p. 868–876.

40. Falzarano, D., O. Krokhin, G. Van Domselaar, K. Wolf, J. Seebach, H.-J. Schnittler and H. Feldmann, Ebola sGP—the first viral glycoprotein shown to be C-mannosylated. Virology, 2007. 368(1): p. 83–90.

41. Plewe, M.B., N.V. Sokolova, V.R. Gantla, E.R. Brown, S. Naik, A. Fetsko, D.D. Lorimer, D.M. Dranow, H. Smutney and J. Bullen, Discovery of Adamantane Carboxamides as Ebola Virus Cell Entry and Glycoprotein Inhibitors. ACS medicinal chemistry letters, 2020. 11(6): p. 1160–1167.

42. Ren, J., Y. Zhao, E.E. Fry and D.I. Stuart, Target identification and mode of action of four chemically divergent drugs against ebolavirus infection. Journal of medicinal chemistry, 2018. 61(3): p. 724–733.

43. Shaikh, F., Y. Zhao, L. Alvarez, M. Iliopoulou, C. Lohans, C.J. Schofield, S. Padilla-Parra, S.W. Siu, E.E. Fry and J. Ren, Structure-based in silico screening identifies a potent ebolavirus inhibitor from a traditional Chinese medicine library. Journal of medicinal chemistry, 2019. 62(6): p. 2928–2937.

44. Zhao, Y., J. Ren, E.E. Fry, J. Xiao, A.R. Townsend and D.I. Stuart, Structures of Ebola virus glycoprotein complexes with tricyclic antidepressant and antipsychotic drugs. Journal of medicinal chemistry, 2018. 61(11): p. 4938–4945.

45. Zhao, Y., J. Ren, K. Harlos, D.M. Jones, A. Zeltina, T.A. Bowden, S. Padilla-Parra, E.E. Fry and D.I. Stuart, Toremifene interacts with and destabilizes the Ebola virus glycoprotein. Nature, 2016. 535(7610): p. 169–172.

46. Cohen-Dvashi, H., M. Zehner, S. Ehrhardt, M. Katz, N. Elad, F. Klein and R. Diskin, Structural basis for a convergent immune response against Ebola Virus. Cell host & microbe, 2020. 27(3): p. 418–427. e4.

47. Dias, J.M., A.I. Kuehne, D.M. Abelson, S. Bale, A.C. Wong, P. Halfmann, M.A. Muhammad, M.L. Fusco, S.E. Zak and E. Kang, A shared structural solution for neutralizing ebolaviruses. Nature structural & molecular biology, 2011. 18(12): p. 1424–1427.

48. Ehrhardt, S.A., M. Zehner, V. Krähling, H. Cohen-Dvashi, C. Kreer, N. Elad, H. Gruell, M.S. Ercanoglu, P. Schommers and L. Gieselmann, Polyclonal and convergent antibody response to Ebola virus vaccine rVSV-ZEBOV. Nature medicine, 2019. 25(10): p. 1589–1600.

49. King, L.B., B.R. West, C.L. Moyer, P. Gilchuk, A. Flyak, P.A. Ilinykh, R. Bombardi, S. Hui, K. Huang and A. Bukreyev, Cross-reactive neutralizing human survivor monoclonal antibody BDBV223 targets the ebolavirus stalk. Nature communications, 2019. 10(1): p. 1–8.

50. Milligan, J.C., C.W. Davis, P.A. Ilinykh, K. Huang, P. Halfmann, R.W. Cross, V. Borisevich, K.N. Agans, J.B. Geisbert and C. Chennareddy, Asymmetric and Non-Stoichiometric Recognition Results in Broad Protection Against Ebolaviruses by a Two-Antibody Cocktail.

51. Misasi, J., M.S. Gilman, M. Kanekiyo, M. Gui, A. Cagigi, S. Mulangu, D. Corti, J.E. Ledgerwood, A. Lanzavecchia and J. Cunningham, Structural and molecular basis for Ebola virus neutralization by protective human antibodies. Science, 2016. 351(6279): p. 1343–1346.

52. Murin, C.D., P. Gilchuk, P.A. Ilinykh, K. Huang, N. Kuzmina, X. Shen, J.F. Bruhn, A.L. Bryan, E. Davidson and B.J. Doranz, Convergence of a common solution for broad ebolavirus neutralization by glycan cap-directed human antibodies. Cell reports, 2021. 35(2): p. 108984.

53. Olal, D., A. Kuehne, S. Bale, P. Halfmann, T. Hashiguchi, M.L. Fusco, J.E. Lee, L.B. King, Y. Kawaoka and J.M. Dye, Structure of an Ebola virus-protective antibody in complex with its mucin-domain linear epitope. Journal of Virology, 2011.

54. Pallesen, J., C.D. Murin, N. De Val, C.A. Cottrell, K.M. Hastie, H.L. Turner, M.L. Fusco, A.I. Flyak, L. Zeitlin and J.E. Crowe, Structures of Ebola virus GP and sGP in complex with therapeutic antibodies. Nature microbiology, 2016. 1(9): p. 1–9.

55. West, B.R., C.L. Moyer, L.B. King, M.L. Fusco, J.C. Milligan, S. Hui and E.O. Saphire, Structural basis of pan-ebolavirus neutralization by a human antibody against a conserved, yet cryptic epitope. MBio, 2018. 9(5): p. e01674–18.

56. Wilson, J.A., M. Hevey, R. Bakken, S. Guest, M. Bray, A.L. Schmaljohn and M.K. Hart, Epitopes involved in antibody-mediated protection from Ebola virus. Science, 2000. 287(5458): p. 1664–1666.

57. Pascal, K.E., D. Dudgeon, J.C. Trefry, M. Anantpadma, Y. Sakurai, C.D. Murin, H.L. Turner, J. Fairhurst, M. Torres and A. Rafique, Development of clinical-stage human monoclonal antibodies that treat advanced Ebola virus disease in nonhuman primates. The Journal of infectious diseases, 2018. 218(suppl_5): p. S612–S626.

58. Bagdonaite, I. and H.H. Wandall, Global aspects of viral glycosylation. Glycobiology, 2018. 28(7): p. 443–467.

59. Hargett, A.A. and M.B. Renfrow, Glycosylation of viral surface proteins probed by mass spectrometry. Current opinion in virology, 2019. 36: p. 56–66.

60. Snijder, J., M.S. Ortego, C. Weidle, A.B. Stuart, M.D. Gray, M.J. McElrath, M. Pancera, D. Veesler and A.T. McGuire, An antibody targeting the fusion machinery neutralizes dual-tropic infection and defines a site of vulnerability on Epstein-Barr virus. Immunity, 2018. 48(4): p. 799–811. e9.

61. Walls, A.C., X. Xiong, Y.-J. Park, M.A. Tortorici, J. Snijder, J. Quispe, E. Cameroni, R. Gopal, M. Dai and A. Lanzavecchia, Unexpected receptor functional mimicry elucidates activation of coronavirus fusion. Cell, 2019. 176(5): p. 1026–1039. e15.

62. Watanabe, Y., J.D. Allen, D. Wrapp, J.S. McLellan and M. Crispin, Site-specific glycan analysis of the SARS-CoV-2 spike. Science, 2020. 369(6501): p. 330–333.

63. Wörner, T.P., T.M. Shamorkina, J. Snijder and A.J. Heck, Mass spectrometry-Based structural virology. Analytical Chemistry, 2020. 93(1): p. 620–640.

64. Yao, H., Y. Song, Y. Chen, N. Wu, J. Xu, C. Sun, J. Zhang, T. Weng, Z. Zhang and Z. Wu, Molecular architecture of the SARS-CoV-2 virus. Cell, 2020. 183(3): p. 730–738. e13.

65. Behrens, A.-J. and M. Crispin, Structural principles controlling HIV envelope glycosylation. Current opinion in structural biology, 2017. 44: p. 125–133.

66. Go, E.P., A. Herschhorn, C. Gu, L. Castillo-Menendez, S. Zhang, Y. Mao, H. Chen, H. Ding, J.K. Wakefield and D. Hua, Comparative analysis of the glycosylation profiles of membrane-anchored HIV-1 envelope glycoprotein trimers and soluble gp140. Journal of virology, 2015. 89(16): p. 8245–8257.

67. Go, E.P., G. Hewawasam, H.-X. Liao, H. Chen, L.-H. Ping, J.A. Anderson, D.C. Hua, B.F. Haynes and H. Desaire, Characterization of glycosylation profiles of HIV-1 transmitted/founder envelopes by mass spectrometry. Journal of virology, 2011. 85(16): p. 8270–8284.

68. Raska, M., K. Takahashi, L. Czernekova, K. Zachova, S. Hall, Z. Moldoveanu, M.C. Elliott, L. Wilson, R. Brown and D. Jancova, Glycosylation patterns of HIV-1 gp120 depend on the type of expressing cells and affect antibody recognition. Journal of Biological Chemistry, 2010. 285(27): p. 20860–20869.

69. Bagdonaite, I., R. Nordén, H.J. Joshi, S.L. King, S.Y. Vakhrushev, S. Olofsson and H.H. Wandall, Global mapping of O-glycosylation of varicella zoster virus, human cytomegalovirus, and Epstein-Barr virus. Journal of Biological Chemistry, 2016. 291(23): p. 12014–12028.

70. Bräutigam, J., A.J. Scheidig and W. Egge-Jacobsen, Mass spectrometric analysis of hepatitis C viral envelope protein E2 reveals extended microheterogeneity of mucin-type O-linked glycosylation. Glycobiology, 2013. 23(4): p. 453–474.

71. Brun, J., S.a. Vasiljevic, B. Gangadharan, M. Hensen, A. V. Chandran, M.L. Hill, J. Kiappes, R.A. Dwek, D.S. Alonzi and W.B. Struwe, Assessing Antigen Structural Integrity through Glycosylation Analysis of the SARS-CoV-2 Viral Spike. ACS central science, 2021. 7(4): p. 586–593.

72. Watanabe, Y., T.A. Bowden, I.A. Wilson and M. Crispin, Exploitation of glycosylation in enveloped virus pathobiology. Biochimica et Biophysica Acta (BBA)-General Subjects, 2019. 1863(10): p. 1480–1497.

73. Larkin, M.A., G. Blackshields, N.P. Brown, R. Chenna, P.A. McGettigan, H. McWilliam, F. Valentin, I.M. Wallace, A. Wilm and R. Lopez, Clustal W and Clustal X version 2.0. bioinformatics, 2007. 23(21): p. 2947–2948.

74. Waterhouse, A., M. Bertoni, S. Bienert, G. Studer, G. Tauriello, R. Gumienny, F.T. Heer, T.A.P. de Beer, C. Rempfer and L. Bordoli, SWISS-MODEL: homology modelling of protein structures and complexes. Nucleic acids research, 2018. 46(W1): p. W296–W303.

75. Goddard, T.D., C.C. Huang, E.C. Meng, E.F. Pettersen, G.S. Couch, J.H. Morris and T.E. Ferrin, UCSF ChimeraX: Meeting modern challenges in visualization and analysis. Protein Science, 2018. 27(1): p. 14–25.

76. Emsley, P. and K. Cowtan, Coot: model-building tools for molecular graphics. Acta crystallographica section D: biological crystallography, 2004. 60(12): p. 2126–2132.

77. de Beer, T., J.F. Vliegenthart, A. Loeffler and J. Hofsteenge, The Hexopyranosyl Residue That Is C-Glycosidically Linked to the Side Chain of Tryptophan-7 in Human RNase Us Is. alpha.-Mannopyranose. Biochemistry, 1995. 34(37): p. 11785–11789.

78. Frank, M., D. Beccati, B.R. Leeflang and J.F. Vliegenthart, C-Mannosylation Enhances the Structural Stability of Human RNase 2. Iscience, 2020. 23(8): p. 101371.

79. Jonker, H.R., K. Saxena, A. Shcherbakova, B. Tiemann, H. Bakker and H. Schwalbe, NMR Spectroscopic Characterization of the C-Mannose Conformation in a Thrombospondin Repeat Using a Selective Labeling Approach. Angewandte Chemie, 2020. 132(46): p. 20840–20846.

80. Bern, M., Y.J. Kil and C. Becker, Byonic: advanced peptide and protein identification software. Current protocols in bioinformatics, 2012. 40(1): p. 13.20. 1–13.20. 14.

81. Pino, L.K., B.C. Searle, J.G. Bollinger, B. Nunn, B. MacLean and M.J. MacCoss, The Skyline ecosystem: Informatics for quantitative mass spectrometry proteomics. Mass spectrometry reviews, 2020. 39(3): p. 229–244.

