## Supplementary Information for "Glycan shield of the ebolavirus envelope glycoprotein GP"

Supplementary Table S1 legend

Supplementary Figures S1-S5

**Supplementary Table S1 legend**

Lists of site-specific N-linked glycan compositions identified with Byonic from LC-MS/MS of EBOV and BDBV GP. Each sample type (HEK293/S2, EBOV/BDBV, GPΔTM/full-length) is provided on a separate tab, with the corresponding sites listed as the header of every column. Provided are the non-redundant lists of glycan compositions from the duplicate experiments, including peptides with missed cleavages. Abbreviations: HexNAc (N-acetylated Glucosamine), Hex (Hexose), Fuc (Fucose), NeuAc (N-acetyl Neuraminic acid).

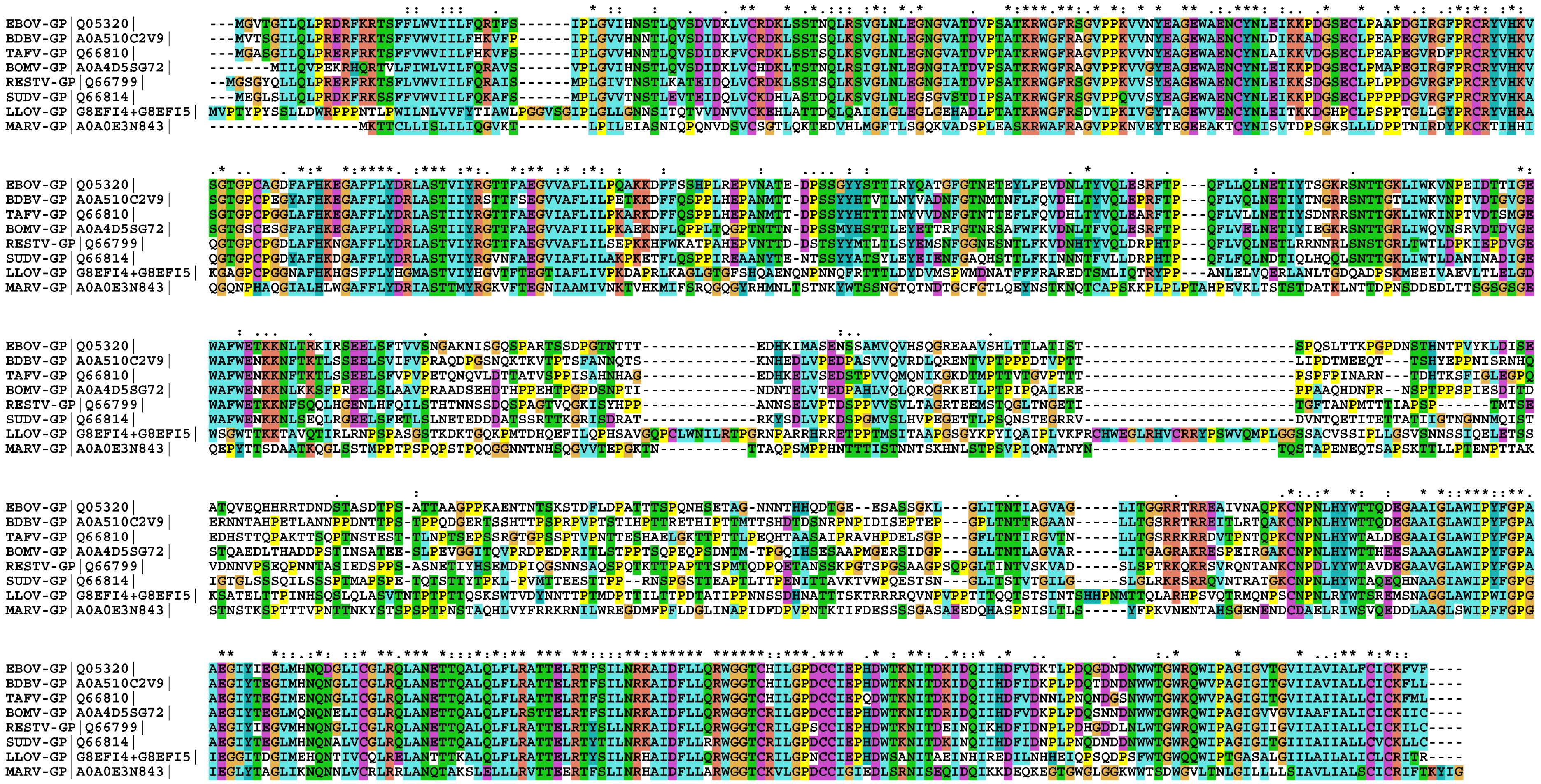


**Supplementary Figure S1.** Multiple sequence alignment of filovirus GP used for the analyses presented in main text (Figure 1). Corresponding UniProt IDs are provided in the sequence names.

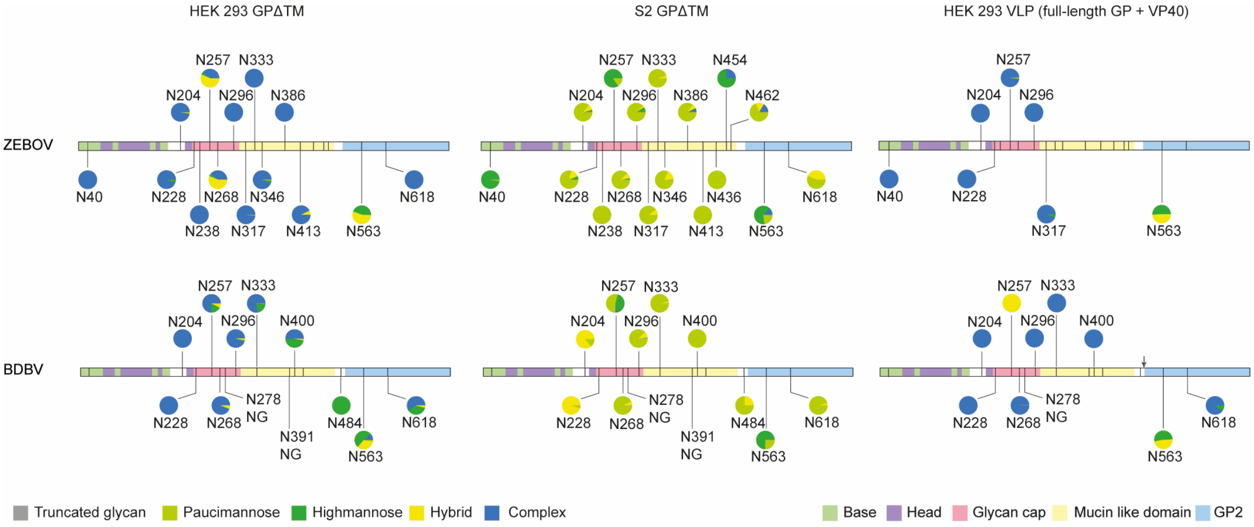


**Supplementary Figure S2.** Overview of site-specific N-linked glycan processing in ebola virus GPΔTM from HEK293 and S2 cells, versus full-length GP from HEK293 derived VLPs as determined by LC-MS/MS. The glycans were classified by HexNAc content as truncated, paucimannose, high-mannose, hybrid or complex. Shown is the average of a duplicate experiment.

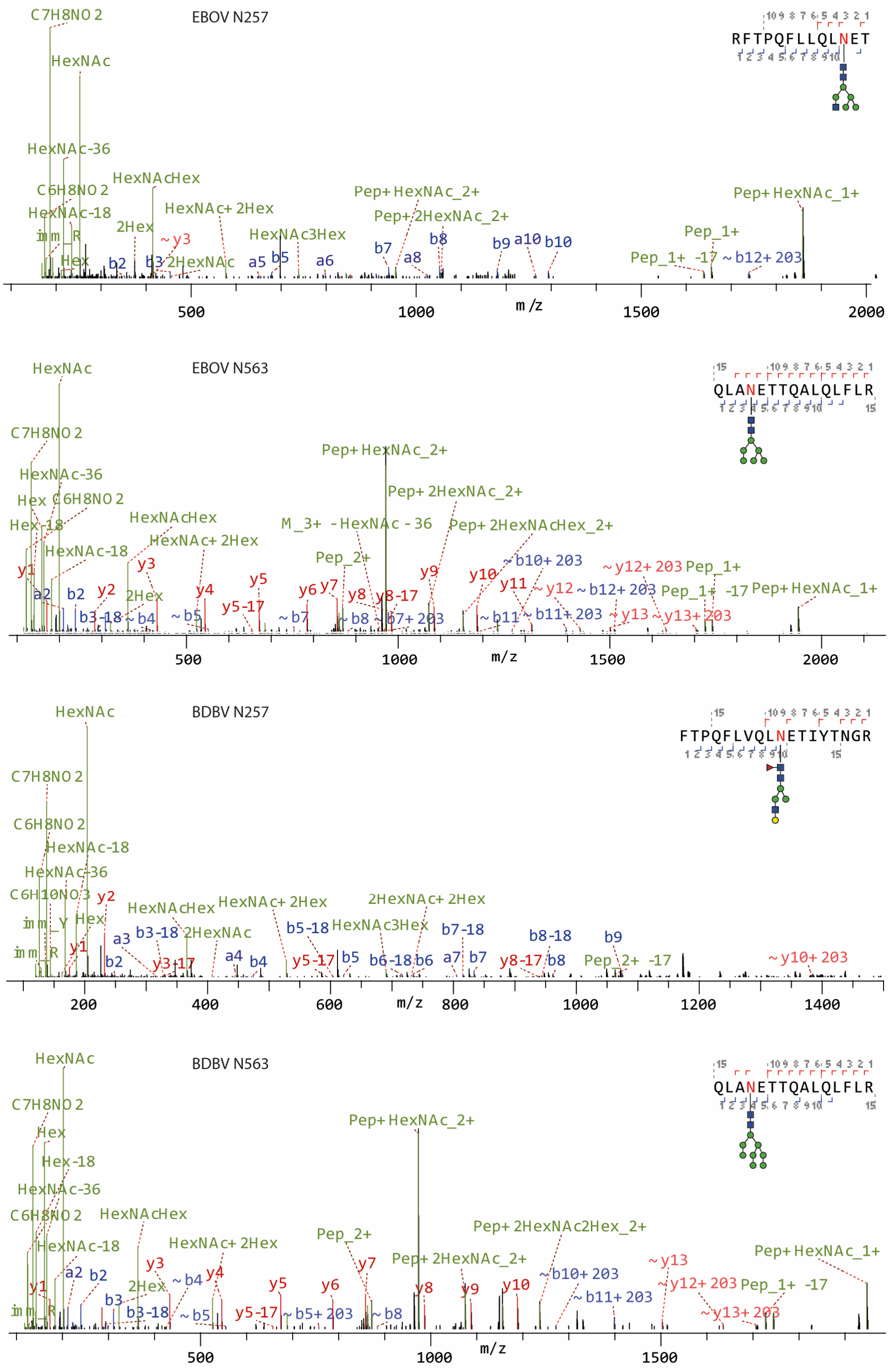


**Supplementary Figure S3.** MS/MS spectra of glycopeptides containing sites N257 or N563 of full-length EBOV and BDBV GP from HEK293 derived VLPs with unprocessed high-mannose and hybrid glycans.

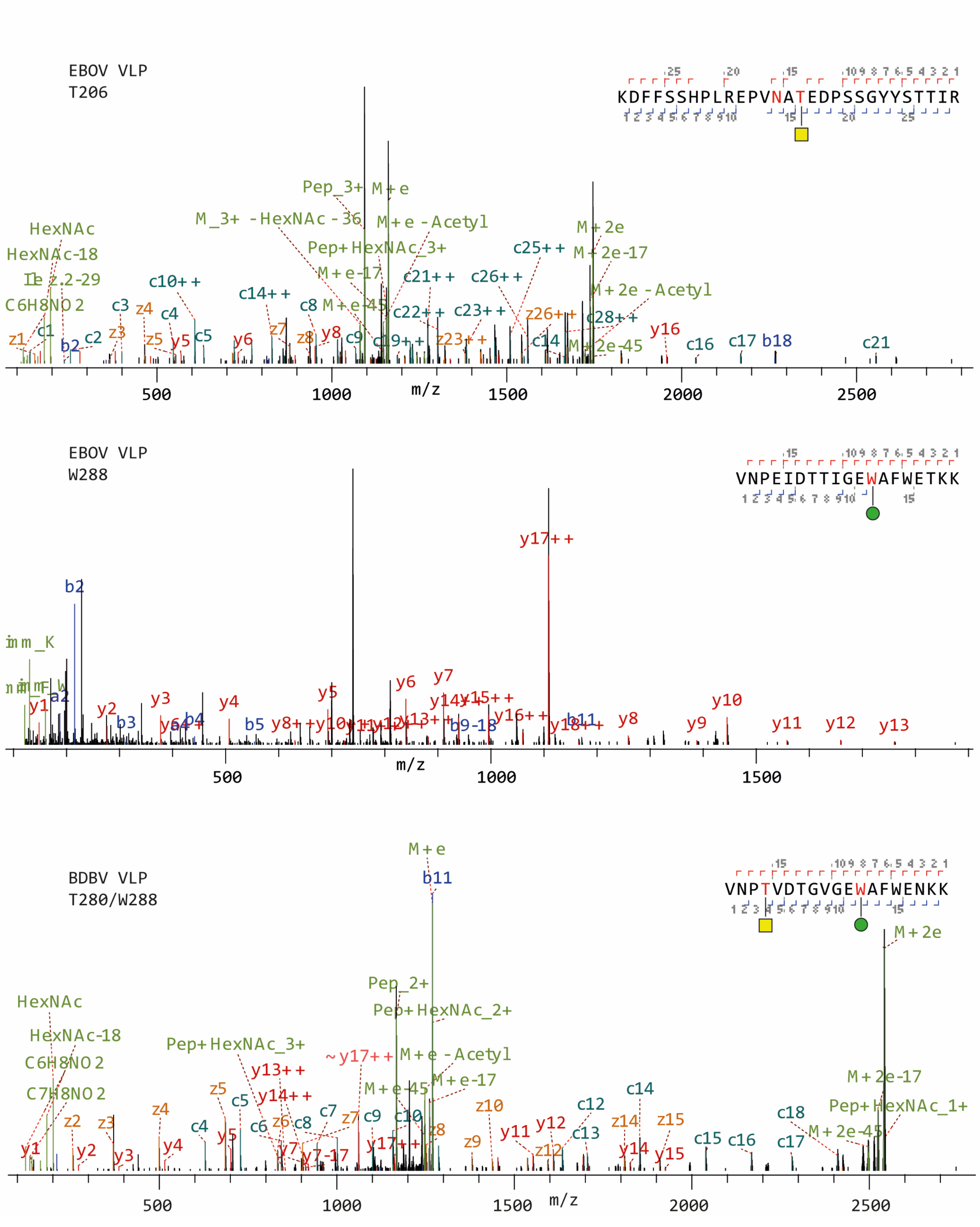


**Supplementary Figure S4.** MS/MS spectra of glycopeptides containing sites O-linked glycosylation outside the MLD and C-linked mannosylation at W288 in full-length EBOV and BDBV GP from HEK293 derived VLPs with unprocessed high-mannose and hybrid glycans.

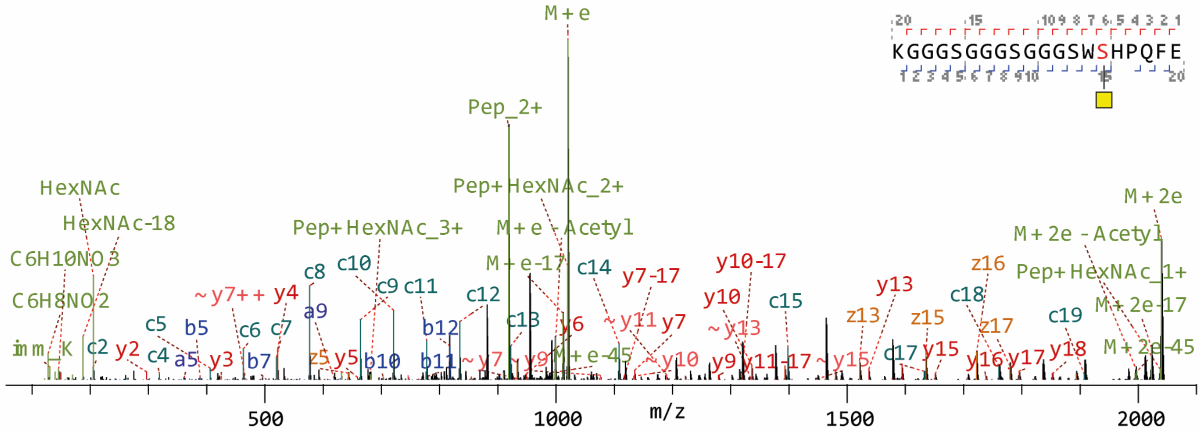


**Supplementary Figure S5.** MS/MS spectrum of glycopeptide from HEK293-derived BDBV GPΔTM with O-linked glycosylation in the Strep-tag of the recombinant soluble ectodomain.
